# Ts-Biotag, a multimodal reporter of Tie2 expression, labels microglia in a model of neuro injury

**DOI:** 10.1101/2023.02.08.527655

**Authors:** Jennifer Brodsky, Zeinab Tashi, Sharon K. Christopher, Gabriella Cerna, Simone Gohsman, Benjamin B. Bartelle

## Abstract

The Ts-Biotag transgenic mouse reports the expression of receptor tyrosine kinase Tie2, a known marker of angiogenic states for both vascular endothelial cells and macrophages. We demonstrate Ts-Biotag labeling and Tie2 expression in a neural injury model to find the majority of labeling occurs in the myeloid derived and brain resident cell type, microglia. Additionally the ligand of Tie2, Ang1, is dynamically expressed, first in astrocytes then neural progenitor during wound signaling and healing. These results offer a Tie2 specific, in vivo view of a neuroimmune response to injury, suggesting a microglia/neural progenitor intercellular interaction guides recovery from a brain lesion.

**Graphical Abstract:** The Ts-Biotag mouse reports expression of Tie2 for any imaging modality compatible with avidinated agents. Mice were given a transcranial cryo-injury and Ts-Biotag activity was followed for 7 days with MRI and histology, showing local and systemic Ts-Biotag labeling. Histology of WT and labeled bone marrow chimeras showed the protein Tie2 expressed in microglia, which assembled at the border of the lesion 1-2d post injury before invading by day 7. The main ligand of Tie2, Ang1, was first expressed systemically by astrocytes, then by neural progenitor cells proximal to and within the lesion. These results elucidate an axis of intercellular signaling involved in the resolution of inflammation and partial healing of a CNS injury.

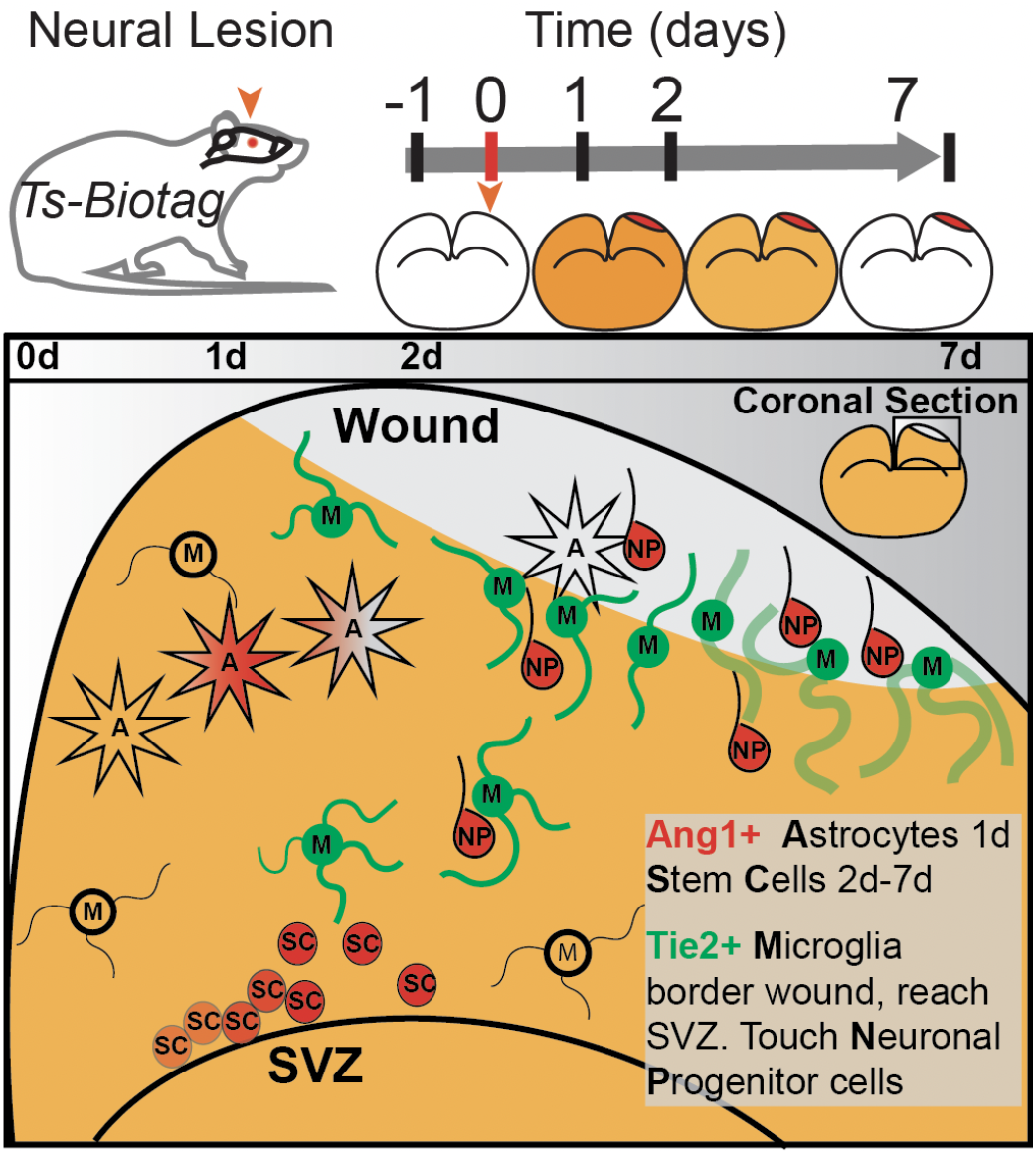

## Introduction

Neural injuries can lead to tissue regeneration or the formation of a non functional scar depending on the organism, aging and the severity of injury. In human pathologies like MS or stroke, a lesion can heal almost completely or become permanently damaged, depending on the wound response mounted by reactive astrocytes and microglia, the innate immune cells of the brain. Understanding the signaling cascades and the cell types involved in wound signaling would offer immediate clinical benefit, but functional and mechanistic studies of these transient events remains challenging. Many key pathways can only work in a living animal, where multiple cell types communicate via paracrine signals. One such axis of tissue-level signaling is the receptor tyrosine kinase, Tie2 and its growth factor ligands, angiopoietin 1 and 2 (Ang1, Ang2).

The biology of the Tie2 receptor tyrosine kinase has been clearly identified in both vascular endothelial cells (VECs)(Hughes et al. 2003) and Tie2-expressing macrophages (TEMs) as mediators of angiogenesis and markers of aggressive tumor growth (De Palma et al. 2005). Conversely, macrophages can secrete Ang1 and Ang2 in response to inflammation signals (Hubbard et al. 2005)(Okuno et al. 2011). The angiopoietins function as proliferative factors in cultured neuronal cells, and immunohistochemistry has shown Ang2 expression in neural progenitor cells (NPCs) in the mouse subventricular zone, one of the few stem cell niches of the adult brain, in response to stroke (Liu et al. 2009).

Microglia, like macrophages, are myeloid derived, with many similar physiological functions, but the role of Tie2 has not been described for this brain resident cell type. The Tie2 gene (TEK) is not expressed in basal adult microglia or in models of MS or AD studied by single cell transcriptomics, (Hammond et al. 2019). TEMs have been identified in brain tumors, however no Tie2+ cells have been identified as microglia (Blank et al. 2021).

We previously developed an in vivo multimodal reporter system driven by the Tie2 promoter enhancer (Ts-Biotag) to look at the expression of Tie2 in vascular development in vivo (Bartelle et al. 2012; Suero-Abreu et al. 2017). In this study, we explore the utility of Ts-Biotag mice to resolve wound responses and healing in a model of neural injury. The in vivo and multimodal imaging capability of the Ts-Biotag system allowed for a broad spatial and temporal view of Tie2 expression in response to injury followed by immunohistology at select time points to more clearly identify Tie2 expressing cells and potential interactions with other cell types. We were surprised to find predominantly myeloid expression of Tie2 and further determined that the cell type was not peripherally derived. Moreover, Ang1 was expressed in both astrocytes and NPC. These findings present potential new biological roles for microglia and intercellular communication via the Ang1/Tie2 signaling axis.

## METHODS

### Cryo-injury

A 4mm blunt probe was maintained at −78°C in dry ice prior to surgery. Mice were anesthetized with isoflurane, secured in a stereotaxic apparatus (**Name of apparatus**), and their heads were shaved. A 2cm incision was made with scissors along the midline of the head and the skin was held apart by blunt curved forceps. The blunt probe was applied directly to the skull for 1 minute and removed to reveal a white patch of frost equal to the probe diameter, confirming contact. The skin was then closed and sealed with vetbond (**Vetbond info**) to prevent opening. Animals were monitored for any signs of discomfort after recovering from anesthesia.

### Administration of Targeted Probes for Imaging

All probes/contrast agents were injected via tail vein using a 27g needle: 150µg (1mg/ml) Av-555 or 10mg (40mg/ml) Av-DTPA-Gd; 1.5µg (15µg/ml).

### In Vivo MRI

Mice were injected with Av-DTPA-Gd 30min prior to anesthesia and imaging. The animals were maintained under anesthesia using isoflurane and magnetic resonance imaging (MRI) data were acquired with a 2D multi-slice T2-weighted fast spin echo sequence to resolve edema (TE/TR = 39/3000ms; in-plane resolution=200µm, slice thickness = 1mm) with a T1-weighted gradient echo sequence to resolve Av-DTPA-Gd enhanced contrast (TE/TR = 4.7/25ms; FA = 90°; 100µm in-plane resolution; slice thickness = 1mm).

### Immunohistochemistry

Immunostaining was performed on cells, fixed on glass coverslips, and on cryo-histological sections (25-μm) acquired from mouse embryos. Staining was accomplished with standard protocols using the following antibodies: rat anti-CD31/PECAM1 (1:100 / BD Biosciences); mouse anti-Iba1 (1:200 / wako); rabbit anti-Tie2 (1:500 / abcam); rabbit anti-Ang1 (1:500 / abcam); and mouse anit-Tau(1/1000 / BD). Secondary antibody staining was performed with Cy3 (red; polyclonal goat anti-rat IgG /Jackson Immunoresearch); Cy2 (green; polyclonal donkey anti-goat IgG / Jackson Immunoresearch); FITC (green; polyclonal goat anti-rabbit IgG / Jackson Immunoresearch). Images were acquired using a Leica DMIRE, CSU10 Yokogawa confocal scanner unit with EM-CCD Hamamatsu digital camera controlled using Volocity 5.2 (Improvision). Images were acquired using a 40X oil lens 1.25NA or a 20X air lens 0.7NA as a stack of 25, 1µm thick slices. Images displayed are maximum intensity projections from full confocal stacks.

### Bone Marrow Chimeras

Bone marrow chimera (BMC) animals were generated according to standard protocols (Holl 2013). Briefly, fasted C57BL/6 mice were given 900Rads from a cesium source. Concurrently ∼8 femurs were collected from the CAG::mRFP1 transgenic mouse (Jackson labs Strain #:**005884)**. Marrow was flushed out using sterile HBSS, then cells were broken up, spun down and resuspended in RPMI. Mature T cells were eliminated from the bone marrow solution, with anti-Thy1/complement, then filtered through a 70 mm mesh and resuspended in RPMI to a concentration of 10 million cells per 150 µl. Cells were injected retro-orbitally into irradiated recipient mice, and monitored daily for 2 weeks then kept for 4 more weeks before experiments.

### Statistical analysis

MRI contrast significance was determined by ANOVA, followed by Tukey ‘s HSD for pairwise analysis. Co-localization in IHC was determined by a pixel-wise Pearson correlation from full confocal microscopy image stacks across multiple images and biological replicates.

## RESULTS

### Ts-Biotag labels Tie2 expression for over 7 days in response to a brain injury

For a longitudinal view of Tie2 activation in an injury model, cohorts of WT and Ts-Biotag animals were imaged over a 9 day time course using MRI (**Figure 1**). At each time point, animals were initially scanned to look for edema and to confirm there was no residual contrast from previous days, then given an IV injection of Avidin-DTPA-Gd to target genetically expressed biotin cell surface tags. WT controls were collected to account for vascular leakage endogenous biotinylation or other potential artifacts of contrast enhanced imaging (**Figure 1A**). Pre-injury Av-DTPA-Gd had no detectable patterns of contrast in WT animals. Post injury, despite injection with Av-DTPA-Gd, contrast was mostly hypointensities around the lesion area, persisting to 7d.

**Figure 1:**
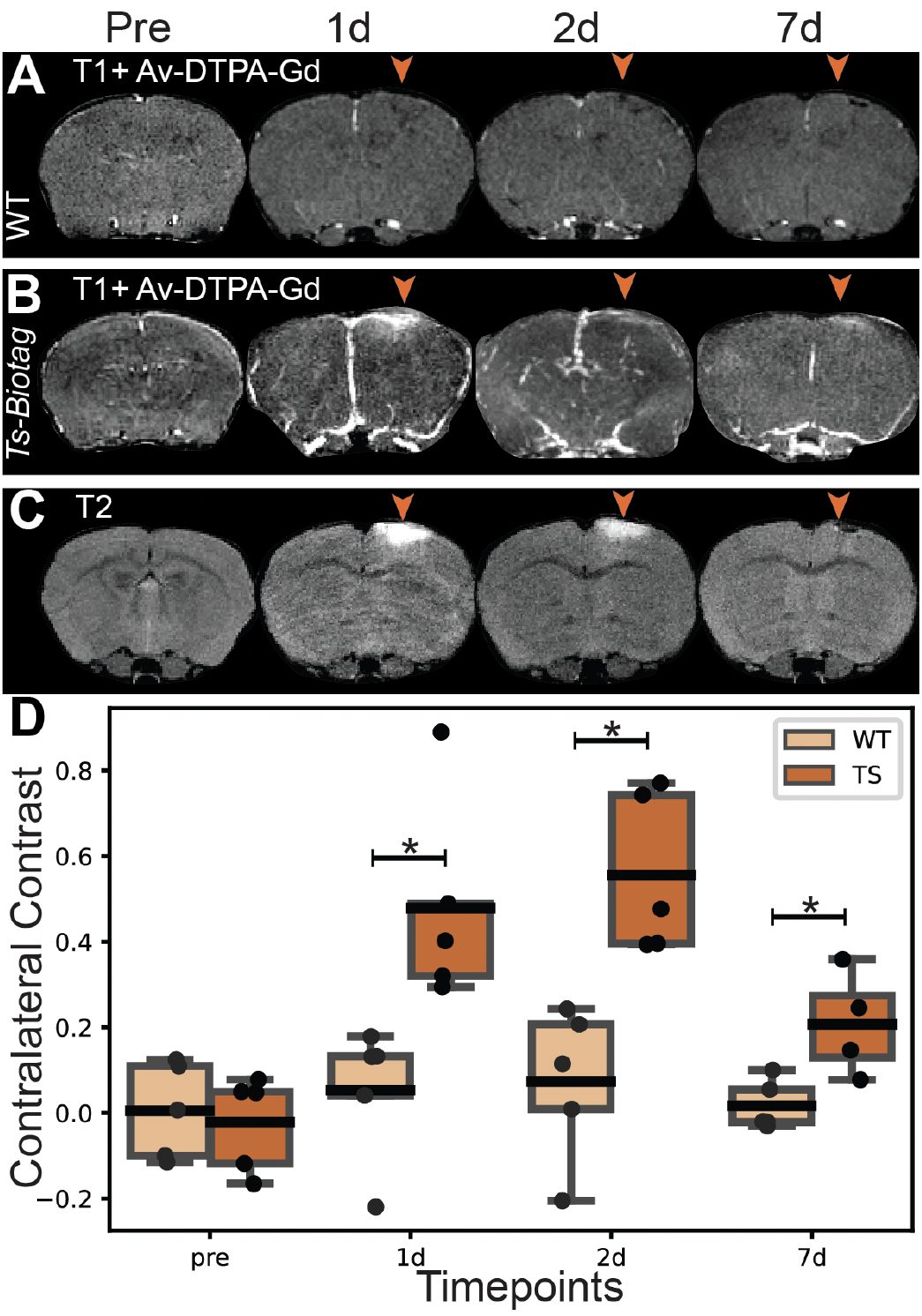
Longitudinal molecular MRI of Ts-Biotag labeling in a cryo-lesion model of neural injury. Longitudinal MRI of WT mouse brain injected IV with Avidin-DTPA-Gd before and after neural injury (**A**). Ts-Biotag animals over the same time course have contrast labeling around the lesion site persisting to 7d post lesion and in larger blood vessels at earlier time points (**B**). Longitudinal T2 MRI shows edema that largely correlates with Ts-Biotag labeling (**C**). Quantitation of Ts-Biotag vs WT contrast over the injury time course. All post injury time points show significant differences (ANOVA/Tukey ‘s p<0.01 n=4) (**D**).

Ts-Biotag animals displayed similar vascular contrast consistent with reported low basal levels of Tie2 (**Figure 1B**). Post injury, Ts-Biotag animals showed bright labeling at the site of the lesion, correlating with edema with contrast enhancement in larger blood vessels. 2d post injury, Ts-Biotag labeling extended to even more vascular. By 7d post injury, no vascular labeling was visible, with only a hyperintense border around the injury site, again coinciding with edema (**Figure 1C**).

Combining longitudinal datasets and single time point experiments, contrast was consistent across replicates, with significant contrast compared to WT controls at each post injury time point (**Figure 1D**). Contrast peaked at ∼50% at 2d post injury, though this sample mean was not significantly higher than that of 1d post injury (p>0.05) even in a paired comparison, demonstrating the high variability of in vivo measurements, even within the same animal.

### Tie2 expression in a cryo-lesion is mostly non vascular

To examine the contribution of specific cell types to overall Ts-Biotag labeling, we generated cryo-lesioned WT mice, harvested brain tissues at the time points used for MRI, and stained sections with immunohistochemistry (IHC). Given the known expression of Tie2 in vascular endothelial cells, we looked for co-localization of Tie2 with the vascular marker CD31.

As with previous studies, adult basal expression of Tie2 was undetectable by IHC (**Figure 2A**). Focusing on tissues immediately adjacent to the cryo-lesion, where Ts-Biotag labeling was highest, Tie2 is highly expressed 1d post injury, however expression was almost entirely non-vascular (**Figure 2B**). Tie2+ cells instead had a “ramified” morphology ascribed to resting state microglia (Lier et al. 2021). Non-vascular Tie2 was abundant at the lesion site 2d post injury, in cells with the more swollen morphology of “activated” microglia (**Figure 2C**). By 7d post injury, CD31 stained cells without vascular morphology were detectable inside of the lesion area (**Figure 2D**). At this late time point, Tie2 staining appeared more diffuse and not clearly associated with one cell type. The lack of clear morphology associated with Tie2 staining at this stage may be due to extracellular cleavage, resulting in “soluble Tie2,” described in the later stages of angiogenesis (Findley et al. 2007).

**Figure 2.**
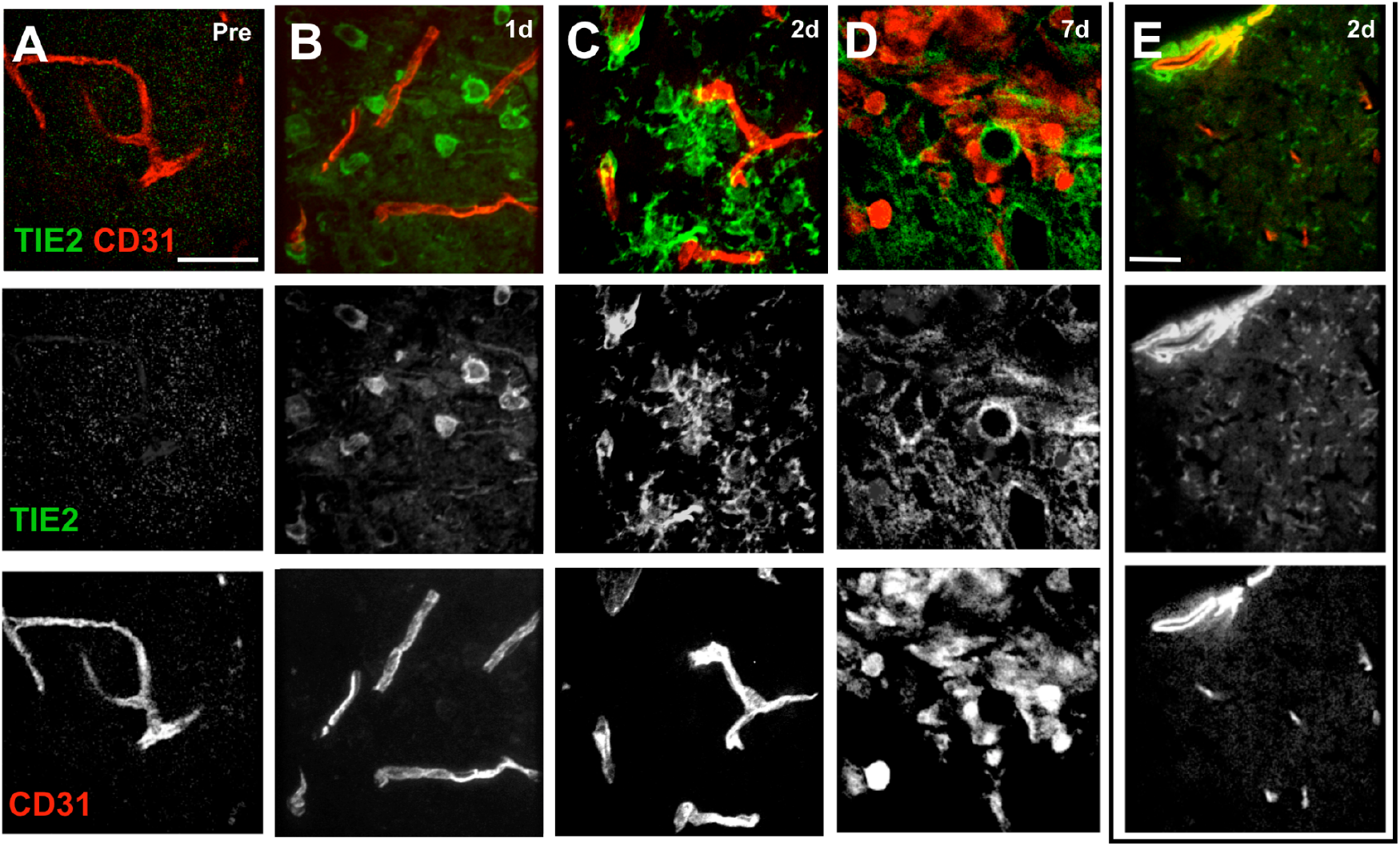
Immunohistology of Tie2 with blood vessel marker CD31 at the border of a neural lesion. Basal levels of Tie2 were not detectable by IHC (**A**). 1d post injury, despite abundant expression, very little Tie2 staining co-localized with blood vessels proximal to the lesion (**B**). 2d post injury, minor vascular labeling persisted with morphological changes in the Tie2+ cell type (**C**). 7d post injury, Tie2 staining lacked cellular morphology and CD31+ cells within the lesion did not appear vascular (**D**) Contralateral to the lesion at 2d post injury, Tie2 was expressed throughout the parenchyma and highly expressed around larger blood vessels (**E**).

Vascular contrast was apparent in MRI at 2d post injury throughout the brain, and looking at the contralateral side of the brain showed colocalization of Tie2 and CD31 in the larger blood vessels of the brain (**Figure 2E**). Tie2 was apparent in fine processes, threaded through the parenchyma throughout the brain, at a lower density than at the wound site, again suggesting microglial expression.

### Tie2 is predominantly expressed in myeloid derived cells in response to a lesion

No one histological marker can specifically identify microglia, but Iba1 is a marker of myeloid derived cells including microglia and macrophages. Prior to injury, microglia had a stereotyped, ramified morphology and remained sparsely tiled across the cortex (**Figure 3A**). 1d post injury, Tie2 expression was detectable mostly in Iba1+ myeloid cells, which increased in density at the edge of the lesion (**Figure 3B**). By 2d post injury, these cells became more compact, but were not the “ameboid” morphology of high inflammation microglia, or invading macrophages (Lier et al. 2021). Diffuse Tie2 marked the edges of the lesion at 7d post injury, with myeloid cells fully invading the lesion and losing most of their ramifications (**Figure 3D**). To further confirm that myeloid cells were both the expressors of Tie2 and our Ts-Biotag reporter, we injected 2d post lesioned mice with Avidin-FITC IV to find uptake of label, almost entirely, by myeloid cells with those proximal to the lesion taking up more label (**Figure 3E**).

**Figure 3.**
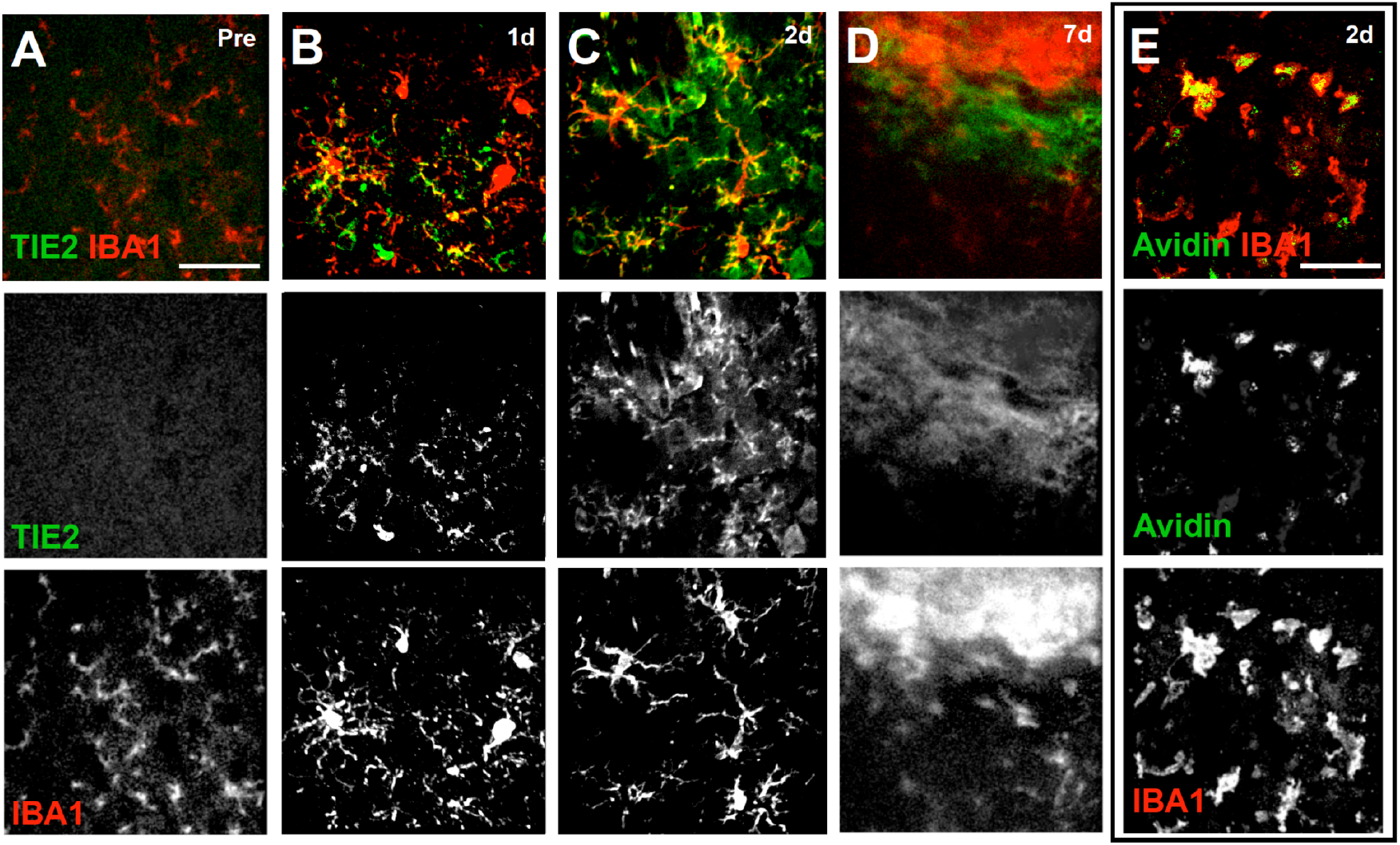
Immunohistology of Tie2 with myeloid marker Iba1 at the border of a neural lesion. Pre-injury, Tie2 was not expressed in Iba1+ cells (**A**). 1d post injury, Tie2 expression bordered the lesion area, with Iba1+ cells at the border and Tie2-cells (**B**). 2d post injury all Iba1+ cells expressed Tie2 (**C**). 7d post injury Tie2 staining was diffuse and lacked morphology with Iba1+ cells both proximal to and invading the lesion site (**D**). Injection of Av-FITC 2d post injury showed labeling almost exclusively in Iba1+ cells proximal to the wound site (**E**).

### Tie2 expressing myeloid cells in a lesion model are not peripherally derived

We sought to determine the identity of the myeloid cells that expressed Tie2 in response to a cryo-lesion model. Circulating macrophages respond to peripheral wound signals by extravasating and invading the site of a lesion, but the blood brain barrier likely modulates this behavior. Without clear histological markers to distinguish microglia from macrophages, we employed BMCs from an Actin-mCherry mouse.

Applying the cryo-lesion to mCherry-BMC mice and harvesting brain tissue at 1-2d post lesion, mCherry+ cells were clearly visible in the lesion site and along the periphery, however these cell types did not have a myeloid morphology and did not co-stain with Tie2 (**Figure 4A**). Instead, mCherry+ cells were highly correlated with the neutrophil marker Ly6G (**Figure 4B**). At 2d post injury mCherry+ cells still did not co-stain with Tie2 (**Figure 4C**), however they were less exclusively neutrophils (**Figure 4D**). Looking specifically for macrophages, Iba1 staining was closely associated with mCherry+ cells, however they were poorly correlated by confocal microscopy (**Figure 4E**). These results suggest that neutrophils are the dominant invasive cell type in this injury model, but other peripheral immune cells do appear, potentially even Tie2-macrophages.

**Figure 4:**
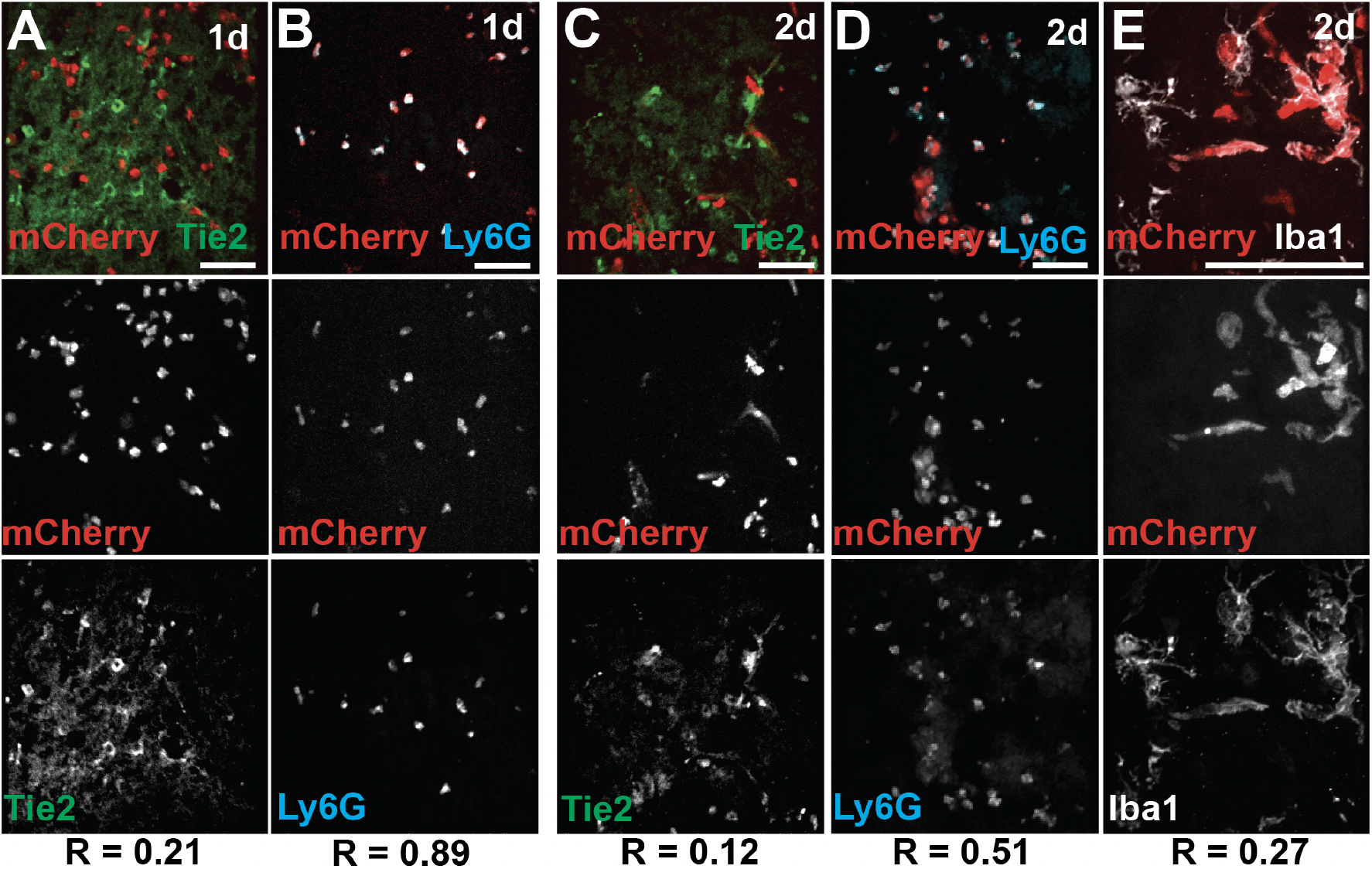
Peripheral cells invading in response to a lesion were Tie2-but express neutrophil markers. IHC of mCherry BMCs, 1-2d post injury. mCherry+ cells appeared proximal to and within the lesion site, but poorly correlated with Tie2 (**A**). The neutrophil marker, Ly6G, marked nearly all mCherry+ cells in the lesion (**B**). 2d post injury mCherry was still not correlated with Tie2 (**C**). Invading cells were no longer entirely Ly6G+ (**D**). Some mCherry+ were Iba1+, but mostly the markers were proximal to each other (**E**).

### Tie2 ligand Ang 1 was first expressed in astrocytes during an early wound response, then by neural progenitor cells

Tie2 expression was upregulated by reactive oxygen species (ROS) and by its own activation by ligand Ang1. Wound signals result in peroxide signaling, but we looked for possible sources of Ang1 that would account for activation of Tie2 over the whole 7d time course.

Ang1 was not detectable by IHC at a basal state in the cortical area targeted for the injury, with some staining around subcortical blood vessels (**Figure 5A**). 1d post injury, Ang1 was highly expressed around the lesion by astrocytes expressing the astroglial marker GFAP (**Figure 5B**). Astrocytic expression of Ang1 was transient, fading at 2d post injury with fewer GFAP+ cells proximal to the lesion at this time point (**Figure 5C**). While cell death within the lesion was occuring, reactive astrocytes did not consistently stain for GFAP and both factors likely contributed to the decline of GFAP+ cells within and proximal to the lesion. More GFAP+ cells became detectable at 7d post injury, suggesting resolution of inflammation (**Figure 5D**). At all later time points, Ang1+ cells were present, but lacked GFAP expression or astrocyte morphology (**Figure 5C,D**).

**Figure 5.**
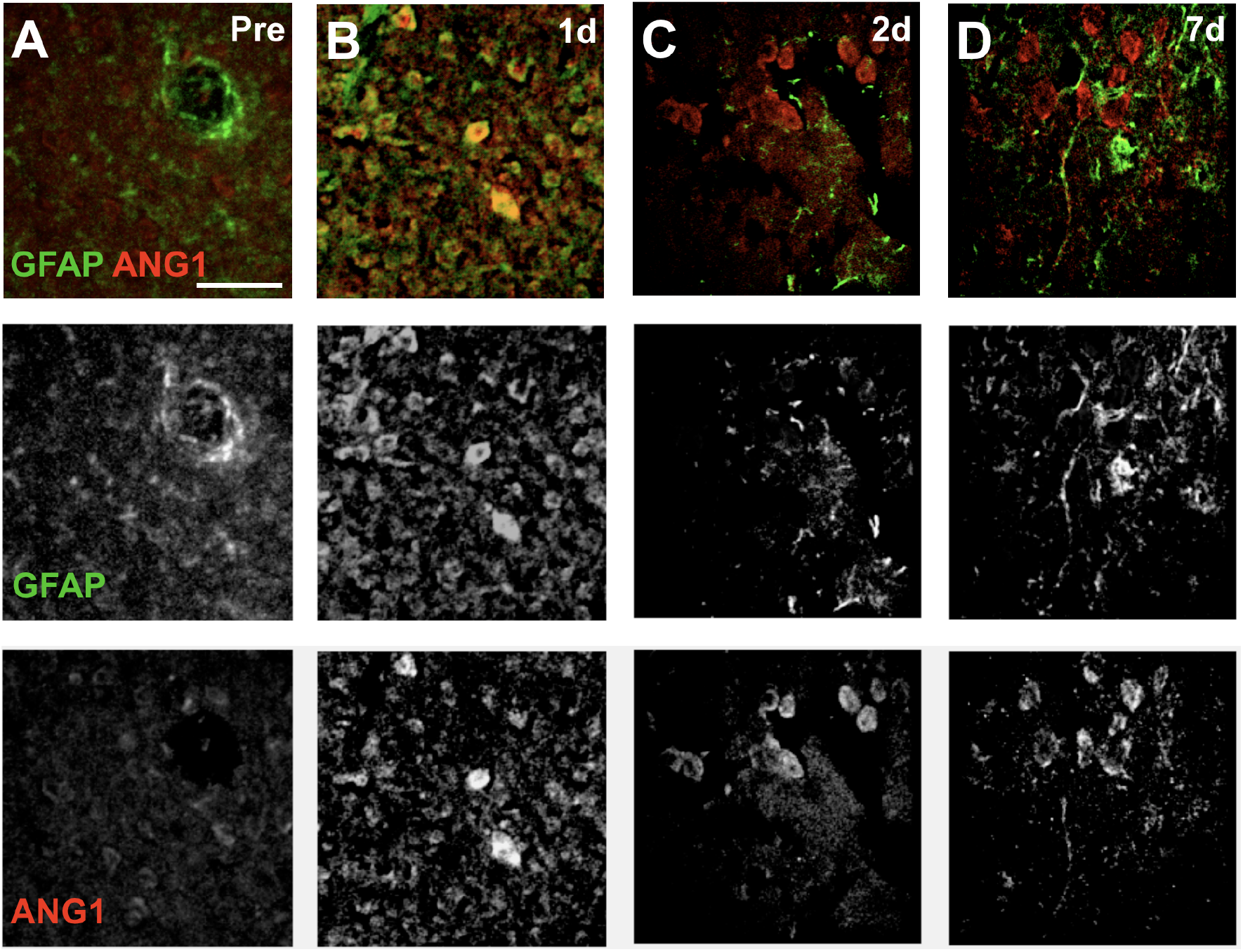
Ang1 was expressed transiently in GFAP+ astrocytes. In a basal brain, GFAP+ cells did not express Ang1 (**A**). 1d post injury, Ang1 broadly co-localized with GFAP+ (**B**). 2d post injury, Ang1 was only expressed in GFAP-cells at the edge of the lesion, with fewer GFAP+ staining in the area (**C**). 7d post injury, more cells were GFAP+ but did not co-stain with Ang1 expression (**D**). Ang1+ cells lacked astrocyte morphology, appearing round, or with a single extending process.

Neural progenitor cells (NPCs) reportedly express Ang1 under some conditions, and 2-7d post injury, the Ang1+ cells proximal to and within the lesion, co-stained with the NPC marker doublecortin (Dcx) (**Figure 6A**). Tie2+ microglia were consistently found proximal to Ang1+ NPCs, spanning from the base of the cortex to the interior of the lesion site (**Figure 6B**). Tie2+ microglia were in partial contact with NPCs, implicating Ang1/Tie2 as an axis of communication between microglia and NPCs in response to injury.

**Figure 6:**
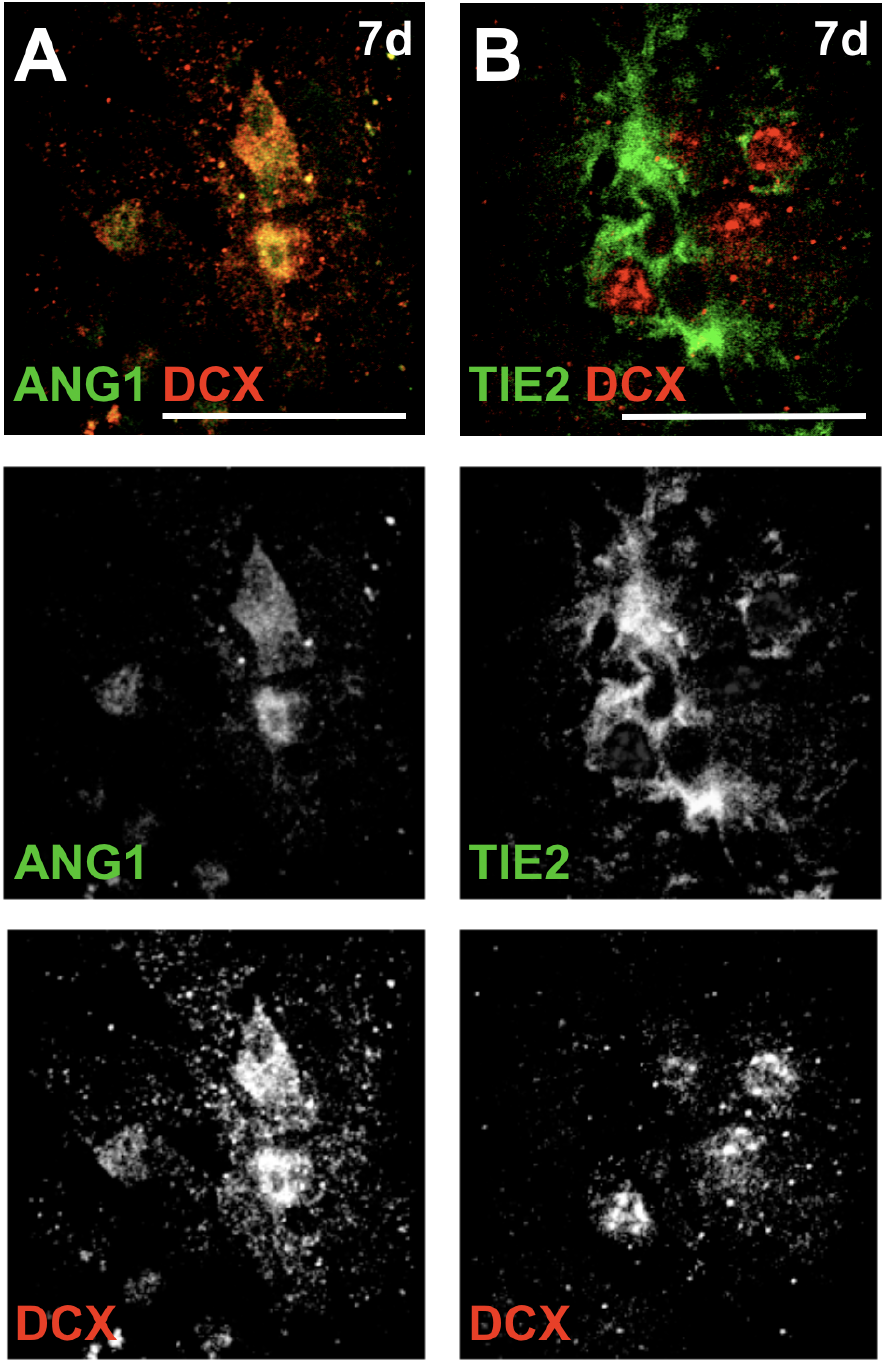
Ang1+ cells were neural progenitors in close contact with microglia. IHC of 7d post injury lesion edge demonstrated Ang1 expression colocalized with neural progenitor cell marker doublecortin (Dcx) (**A**). Dcx+ cells appeared proximal to Tie2+ microglia in this region (**B**).

## DISCUSSION

In this study, a cryo-injury was applied to the Ts-Biotag animal to report Tie2 expression in response to a neural lesion. Within this model, Biotag was accessible to labeling by IV injection, allowing for selective contrast in transgenics that correlated with Tie2 expression. This was almost entirely in myeloid cells and BMC experiments eliminated the possibility that these cells were peripherally derived macrophages. The Tie2 ligand, Ang1 was also highly expressed in response to the cryo-injury, first both broadly and transiently by GFAP+ astrocytes 1d post injury, then by Dcx+ NPCs proximal to the lesion site for at least another 6d.

Microglial access to neurovasculature, as evidenced by robust labeling of Iba1+ cells in Ts-Biotag animals (**Figure 3E**), was unexpected. The Ts-Biotag animal is currently the only example of its application in vivo, but all controls indicate that avidinated agents are limited to the vasculature (Bartelle et al. 2012; Suero-Abreu et al. 2017). Ts-Biotag labeling proximal to the lesion was concurrent with, and mostly limited to areas with edema. Vascular Ts-Biotag labeling was visible 2d post injury across the brain in larger blood vessels, but microglial expression of Tie2 at that time point was detectable by IHC brainwide throughout the parenchyma (**Figure 1C,3E**). Based on these data, both Ts-Biotag expression and edema are necessary for in vivo Biotag labeling of microglia and this constraint must be considered in the use of Biotag as a reporter for in vivo imaging of gene expression in microglia. Tie2 expression, however, is separate from the neuroinflammatory mechanisms that cause edema (Tanaka et al. 2018; Tanaka et al. 2021).

To our knowledge, microglial specific Tie2 expression has not been previously reported, but the gene is well described in macrophages, both during development and by tumor associated macrophages (TAMs) as a marker of rapid angiogenesis and poor prognosis in cancer (Pucci et al. 2009). Peripheral macrophages are thought to cross the BBB in response to inflammatory signaling and may invade an aggressive brain tumor with disrupted vasculature (Liu et al. 2010), though we saw no evidence of macrophage invasion in this injury model.

The dynamic expression of Tie2 and Ang1 goes from an early systemic signal, propagated by astrocytes, that settles into a defined wound response program, reinforced by NPCs. Both receptor and ligand are detectable in proximal cells at all post injury timepoints, implying a signaling axis between microglia and their partner cell types. In culture, Tie2/Ang1 signaling has been characterized as an activator of the AKT/mTOR pathway, leading to increased motility and survival (Bai et al. 2009). If Tie2 is acting similarly in vivo, it could be part of the mechanism that localizes microglia around the lesion area, or facilitates pro-genic signaling between microglia and the invading stem cells.

Tie2 signaling engages myeloid cells in the patterning of tissues developmentally, with vascular tissues best described (Pucci et al. 2009; Nucera et al. 2011). Although it is known that these same pathways can activate in adulthood pathologically during cancer, the Ts-Biotag animal has demonstrated that the same developmental mechanisms can activate as part of a wound response and subsequent healing of a neural lesion. Ang1, as secreted by NPCs, has been previously described developmentally exclusively in relation to vascular endothelial cells as a source of differentiation signaling (Paredes et al. 2021). Whether microglia serve this role in adult tissues and the developmental fate of Ang1+ NPC ‘s in an adult lesion model needs to be determined.

## Abbreviations

Ang1: Angiopoietin 1:
Ang2: Angiopoietin 2:
BMCs: bone marrow chimeras:
Dcx: doublecortin:
IHC: immunohistochemistry:
MRI: magnetic resonance imaging:
NPCs: neural progenitor cells:
ROS: reactive oxygen species:
TAMs: tumor associated macrophages:
TEMs: Tie2-expressing macrophages:
VECs: vascular endothelial cells:

## Acknowledgements

The authors would like to thank the following people: Daniel H Turnbull for guidance and use of his MRI system for the study. Dr Miyeko Mana for use of her confocal and advice on microscopy. Dr Wenbio Gan for his kind gift of the CAG::mRFP1 transgenic mice, and Dr Christopher Parkhurst for instruction in generating BMC mice and guidance in experimental design.

## Attribution

**BB** designed all experiments and wrote the manuscript. **JB** applied the injury model and performed mouse colony management. **ZT** and **SKC** performed MRI. **GC** edited the manuscript. **SG** wrote statistical analysis scripts of imaging data.

## Notes

### Competing Interest Statement

The authors have declared no competing interest.

